# Rapid TetOn-mediated gene expression in neurons across the lifespan with uTTOP

**DOI:** 10.1101/2023.12.05.570307

**Authors:** Christopher K Salmon, Gael Quesseveur, Gee Hung Leo Cheong, Sabrina Chierzi, Miranda Green, Michael P Rosen, Andy YL Gao, J Benjamin Kacerovsky, Keith K Murai

## Abstract

Conditional expression of genes of interest is essential for interrogation of cellular development and function. Although tools exist for conditional gene expression, techniques for rapid-onset, temporally precise expression are lacking. The doxycycline-inducible TetOn expression system allows for this in numerous organ systems, however, transgenic TetOn expression cassettes become silenced in the nervous system during postnatal development. Here, we circumvent this silencing with uTTOP: *in utero* electroporation of Transposable TetOn Plasmids. When electroporated as transposable elements that integrate into the genome, the TetOn system allowed for robust DOX-dependent induction of expression across the postnatal lifespan of the mouse. We demonstrated induction in neurons of sensorimotor and retrospleninal cortex, hippocampus and the olfactory bulb. Latency to peak induction was ≤12 hours, a several fold increase in induction kinetics over existing methodology for *in vivo* conditional expression. To demonstrate the utility of uTTOP, we induced ectopic expression of Sonic hedgehog in adult mouse layer 2/3 cortical neurons, demonstrating that its expression can diversify expression of Kir4.1 in surrounding astrocytes. The rapid induction kinetics of uTTOP allowed us to show that Kir4.1 upregulation significantly lags onset of Shh expression by ∼2 days, a difference in expression time course that is likely not resolvable with current methods. Together, these data demonstrate that uTTOP is a powerful and flexible system for conditional gene expression in multiple brain areas across the mouse lifespan.

## Introduction

Obtaining control over the timing of expression and silencing of genes of interest has been a central challenge in molecular biology. While transgenic approaches have proven to be critical in determining gene function though expression of exogenous molecules or disrupting endogenous gene function, this classical methodology is hampered by the fact that genes play multiple roles across cell types and at different stages of the life cycle of cells and organisms (Luo et al., 2018). For this reason, identification and validation of molecular and genetic switches for controlling gene expression *in vivo* have long been pursued (Lewandoski, 2002). Identifying fast-acting and robust conditional gene expression systems is of particular importance in neurobiology as neural circuit development involves many complex and temporally constrained steps that occur in rapid succession, and genes that regulate development perform diverse functions in the adult as well. While mouse lines expressing variants of Cre-, Dre- and Flp-recombinases are available for temporal control of gene expression in vivo, there remain limitations in these approaches for understanding the function of molecules and pathways in neurons and glial cells of the CNS. For tamoxifen-inducible variants such as CreER/CreERT2 and FLPeR/FlpERT2, high recombination efficiency through tamoxifen-based induction can be challenging especially in adult mice where recombination efficiency can be low (Farmer et al., 2016; Slezak et al., 2007). Cre-, Dre- and Flp-based techniques also lack the ability to titrate gene expression levels as the transcription of the gene-of-interest relies on recombination and upstream promoter activity, and not dosage of a drug or transcription factor. These approaches generally rely on invasive methodology, such as craniotomy and viral injection, as well as fibre optic implantation for photo-activatable recombinases (Meador et al., 2019). Finally, while attempts have been made to enhance induction speed in these systems, the shortest reported latencies to peak expression are still ∼50-72 hours (Sando et al., 2012; Yokoi et al., 2007). Thus, additional approaches are needed to overcome these limitations and to complement the toolkit currently provided by existing methodology.

The TetOn and TetOff tetracycline transactivation systems have been successfully employed to control the timing of gene expression in numerous organ systems, as well as in a wide array of in vitro contexts for probing cell signalling (Baron and Bujard, 2000; Benedetti et al., 2022). The TetOff expression system employs the tetracycline transactivator (tTA), which binds to the tet operator (*tetO*) in the absence of substrate (tetracycline or doxycycline), thereby driving gene expression. TetOn employs the reverse tTA (rtTA), which drives expression in the presence of substrate. Both TetOn and TetOff expression systems have been employed in multiple cell types and regions in the nervous system (Agulhon et al., 2010; Mayford et al., 1996). However, while the TetOff system has been used frequently for inducible gene expression in the CNS (Luo et al., 2018), it has been found that the TetOn system is progressively silenced in the adult mouse CNS, and as a result it has been seldom used (Zhu et al., 2007). Importantly, the TetOff system requires that mice be reared on doxycycline (DOX) diets to supress gene expression until it is desired. DOX is lipophilic and long-term administration leads to its deposition in muscle and bone, meaning that induction upon removal of the DOX diet takes up to 20 days (Mansuy and Bujard, 2000). On the other hand, TetOn-mediated gene expression is observed as few as 4 hours after DOX administration begins (Kistner et al., 1996; Watanabe et al., 2007). With these rapid induction kinetics in mind, the TetOn system has major untapped potential for studying gene function in the nervous system.

Since its initial publication, *in utero* electroporation (IUE) has proven to be a flexible tool for gene expression in neurons *in vivo* (Fukuchi-Shimogori and Grove, 2001; Saito and Nakatsuji, 2001; Tabata and Nakajima, 2001). IUE is a powerful and flexible tool for genetically manipulating neurons with several advantages over other approaches: genetic cassette size is not a limiting factor as it is for viral transduction of genes of interest; new gainor loss-of-function constructs can be tested within days with IUE, rather than months or years as is the case with creation of transgenic animals; IUE is affordable, requiring only plasmid DNA, animals and an electroporator. The introduction of oriented bipolar electric fields and triple-electrode approaches has also made it possible to specifically target numerous brain areas using IUE (Baumgart and Baumgart, 2016; Carlson et al., 2011; dal Maschio et al., 2012). For these reasons, IUE remains a widely used technique for interrogating fundamental cell biology (Bandler et al., 2022; Chung et al., 2023; Ellender et al., 2019; Felske et al., 2023; Gonçalves et al., 2020; Huszár et al., 2022; Mavrovic et al., 2020; Paolino et al., 2023; Prieur et al., 2023; Tsai et al., 2020) and disease (Bennison et al., 2023; Braz et al., 2022; Cases-Cunillera et al., 2021; Druart et al., 2021; Sun et al., 2021) in the brain.

Given the flexibility of IUE for targeting distinct neuronal populations in the CNS and the possibility of achieving fast induction and on-off control of gene expression using TetOn expression systems, we sought to combine these techniques. When introduced by IUE, standard, episomal TetOn plasmids were silenced by postnatal day 21 (P21). However, introducing the TetOn system in piggyBAC transposable elements conferred rapid onset conditional gene expression in multiple brain regions from P21 to 20 months. Latency to peak expression using was ≤12h. We refer to this methodology as uTTOP: *in utero* electroporation of Transposable TetOn Plasmids. Using uTTOP, we achieved conditional gene expression in multiple cortical areas, hippocam-pus and olfactory bulbs. As a proof-of-concept for the utility of uTTOP, we demonstrated that the potent developmental morphogen, Sonic Hedgehog (Shh), can be conditionally expressed in adult neurons to non-cell autonomously alter the molecular properties of neighboring astrocytes in the mature mouse brain. Thus, our results suggest that uTTOP is a powerful tool for conditional expression of genes of interest across the lifespan in mouse neurons.

## Results

### TetOn System Induces Robust Gene Expression in Culture

To assess the utility of the TetOn system in the IUE context, we tested plasmids coding for an enhanced reverse tetracycline transactivator (rtTA2^*S*^-M2, referred to hereafter as rtTA) and a TetOn-inducible ZsGreen1 (ZsG)(Matz et al., 1999). In the presence of doxycycline (DOX), rtTA binds to the tet-operator (tetO) upstream of ZsG in the TetOn plasmid (pTetO-ZsG), thereby inducing expression of ZsG. ZsG is useful for sensitive detection of expression as its quantum yield is much higher than that of EGFP (Kaishima et al., 2016) and it forms aggregates that are more clearly visible than diffuse fluorescent proteins at low expression levels.

We first assessed the induction potential of this system by coexpressing pCMV-rtTA and pTetO-ZsG in HEK293T cells and found that both 0.1µg/mL and 1µg/mL DOX induced ZsG expression approximately two orders of magnitude over control levels (Figure S1A-C). The number of cells displaying leaky expression was low but apparent, however this was likely caused in part by amplification of the pTetO-ZsG plasmid, which contains the SV40 origin of replication, by the large T antigen expressed in HEK293T cells. We next tested induction of the TetOn system in primary cultures of hippocampal neurons. To restrict expression of rtTA to neurons, and to avoid the regulation of the CMV promoter by neuronal depolarization (Wheeler and Cooper, 2001), we replaced the CMV promoter with a human Synapsin1 promoter fragment (hSyn) to produce pSyn-rtTA, which has been used extensively for targeting viral transduction specifically to neurons (Kügler et al., 2003; Rincon et al., 2018). Furthermore, hSyn drives only middling levels of expression from episomal plasmids, which has the added benefit of mitigating leaky induction and transcriptional squelching by high levels of rtTA (Baron and Bujard, 2000). We co-transfected dissociated neuronal cultures with pSyn-rtTA, pTetO-ZsG and a plasmid encoding constitutively expressed tdTomato (pCA-tdTom) as a positive transfection marker. Induction with 0.1 and 1 µg/mL DOX resulted in robust ZsG expression (Figure S1D,E). Control conditions showed very low background green fluorescence.

### Episomal TetOn System Induces Expression in Neurons *in vivo* During Early Development but Not at More Mature Time Points

We next tested DOX-mediated expression of ZsG in neurons *in vivo*. We introduced the same episomal pCAG-rtTA and pTetO-ZsG into layer 2/3 somatosensory cortical neurons at equal concentrations of 1µg/mL via IUE at embryonic day 15-16 (E15-16). We included pCA-tdTom (0.5 µg/mL) as a positive control for electroporation. To avoid invasive injections in P6 pups, we administered DOX to the mother in food (3 mg/g body weight) and via daily IP injections (50µg/g per day) to test whether DOX could be delivered through the milk. This caused robust ZsG expression in somatosensory cortex of P8 pups following 48 hours of administration, demonstrating that DOX-mediated induction can be attained without physical interventions on electroporated pups (Figure 1B).

**Figure 1.**
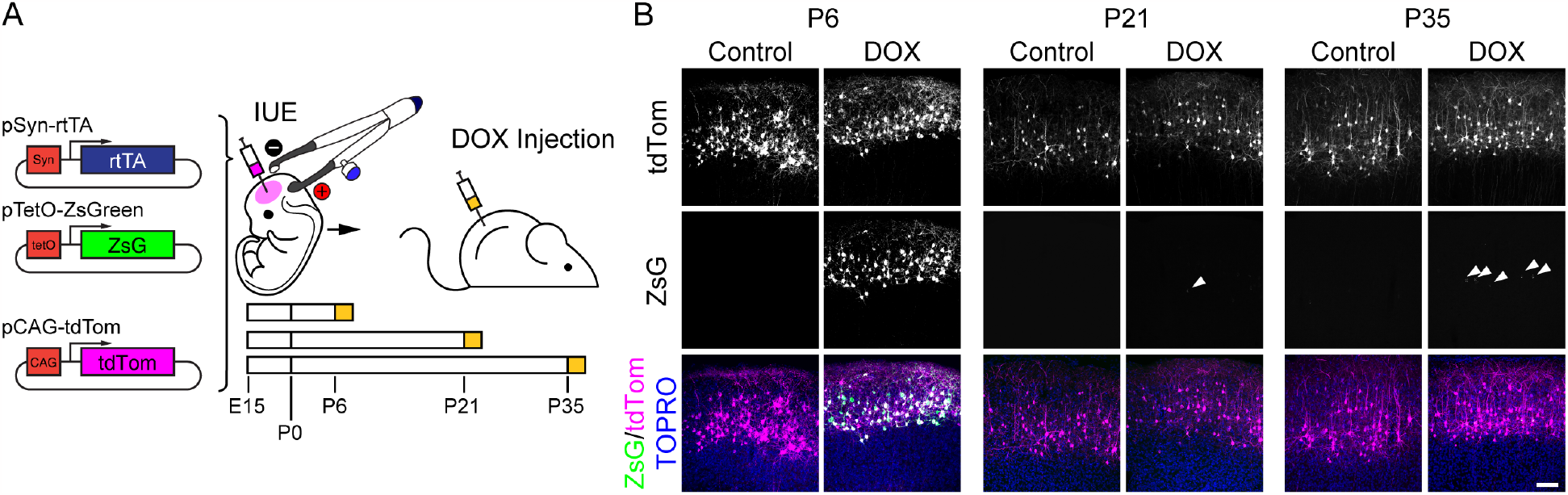
Episomal TetOn allows for induction in cultured neurons and postnatal neurons *in vivo*. **A)** Left, schematic of plasmids for testing DOX-inducible expression of ZsG in the mouse brain following IUE. Reverse tetracycline transactivator (rtTA, purple), driven by the human synapsin promoter fragment (hSyn), activates ZsGreen (ZsG, green) expression via the Tet-response element-based Tet operator (TetO). tdTomato (tdTom), is expressed constitutively from a CAG promoter as a positive control. Right, schematic of IUE (including intrahemispheric injection and tweezer electrodes) and subsequent DOX induction (top) and the three timecourses of DOX induction used in B (bottom). **B)** Induction of ZsG following two days of DOX administration via the mother’s milk is possible at P6 (left), but DOX does not induce strong ZsG expression at P21 (centre) or P35 (right). White arrow heads denote cells displaying low levels of expression. Scale bar 100µm (B).

We next investigated the potential for DOX-mediated induction in P21 and P35 mice that had been electroporated with pSyn-rtTA, pTetO-ZsG and pCA-tdTom (Figure 1A). No ZsG expression was observed following 3 days of DOX administration at P21 (two IP injections of 50µg/g, daily) (1 µg/µL pTetO-ZsG, Figure 1B; 0.1 µg/µL pTetO-ZsG, Figure S2A). Induction was absent at P35 as well. Interestingly, in an experiment using C57B/6 mice, rather than CD1 mice, induction at P21 yielded sparse, low level ZsG expression (Figure S2), suggesting species-specific variation in silencing of the TRE. Thus, while IUE of episomal TetOn plasmids allows for induction at one week of age, this system is not useful for expression at 3 weeks or later. This result is expected from previous work suggesting that tetO-based promoters are permanently silenced in neurons during development, whether present as a transgene or episomally via viral transduction (Zhu et al., 2007). Enhanced delivery of DOX across the blood brain barrier did not improve induction, suggesting that optimization of DOX administration would not greatly improve induction in electroporated neurons. An alternative approach is therefore needed to circumvent tetO silencing.

### Robust DOX-Mediated Induction with uTTOP

An important caveat in the silencing of the TetOn system is that some transgenic mice harbor neuron-specific TetOn expression cassettes that allow for induction (Mansuy et al., 1998; Yamamoto et al., 2003), while others have little or no penetrance (Beard et al., 2006; Eckenstein et al., 2006; Uchida et al., 2006; Zhu et al., 2007). This suggests that the integration site of tetO-controlled constructs may modulate the extent of induction. We therefore hypothesized that the semi-stochastic genomic integration provided by the piggyBAC (PB) transposable expression system (Ding et al., 2005) would overcome TetOn silencing observed in other studies. In the presence of piggyBac transposase derived from the moth, *Trichoplusia ni*, inverted terminal repeats (ITRs) that flank the transposable element (i.e. the gene of interest) allow for homologous recombination of the construct into the genome at TTAA tetranucleotide sequences (Fraser et al., 1995, 1996; Toshiki et al., 2000). This system has been used previously in conjunction with IUE to allow for genomic integration of genetic constructs in astrocytes which, due to the postnatal proliferation of their progenitors, are difficult to target with IUE of episomal plasmids (Figueres-Oate et al., 2016; García-Marqués and López-Mascaraque, 2013). We therefore cloned the promoters and coding sequences of pTetO-ZsG and pSyn-rtTA into piggyBac (PB) transposable elements, creating the uTTOP expression system (*in utero* electroporation of Transposable TetOn Plasmids).

We targeted uTTOP plasmids (PB-pTetO-ZsG, PBpSyn-rtTA) to layer 2/3 somatosensory cortical neurons by IUE at E15-16 (Fig. 2A), electroporating 1µg/µL PBpSyn-rtTA with 0.1 or 1 µg/uL PB-pTetO-ZsG. We induced expression at P35 for 2 days (two IP injections of 50µg/g DOX per day). In both cases, we observed robust induction of ZsG that spanned the entire field of electroporated cells marked by constitutive tdTom expression (Figures 2B, S3). The majority of electroporated cells in DOX-treated animals coexpressed ZsG and tdTom (Fig. 2B and S3).

**Figure 2.**
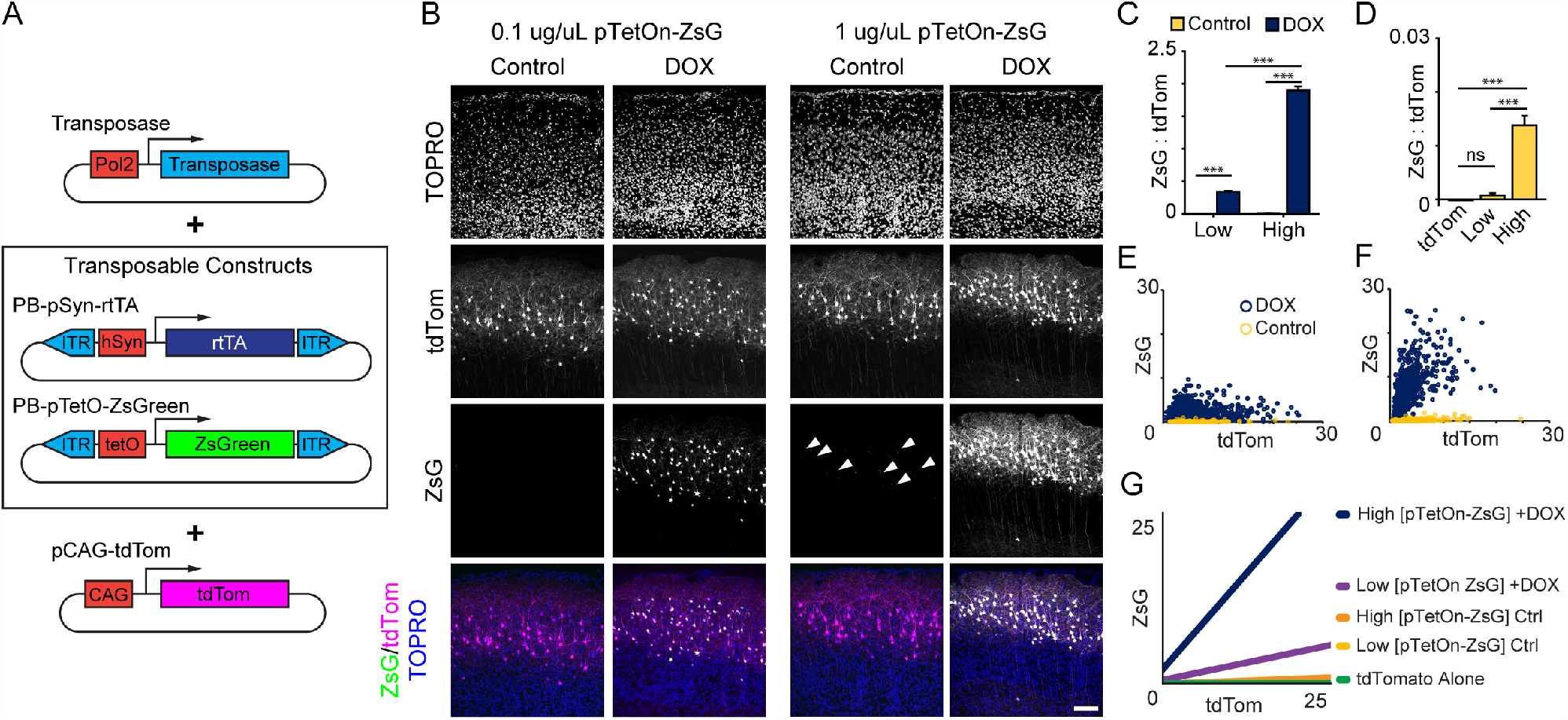
uTTOP allows for robust induction in adult neurons with little or no detectable leak. **A)** Schematic of plasmids used for testing DOX-inducible expression of ZsG by uTTOP. **B)** Images of epicentre of electroporated cortical field from animals electroporated with 0.1 (Low, left) or 1 (High, right) µg/µL PB-pTetO-ZsG, respectively. White arrow heads denote cells displaying low levels of leaky expression. **C)** Plot of mean corrected total cell fluorescence (CTCF) green fluorescence normalized to red CTCF to take into account variable efficiency of electroporation from cell to cell (Low Ctrl 8.06x10-4±0.03; Low DOX 0.34±0.02; High Ctrl 0.014±0.04; High DOX 1.90±0.03; 2-way ANOVA p<0.001 for Treatment, Condition and Interaction.) **D)** Comparison of green fluorescence from uninduced Control animals electroporated with tdTom alone, 0.1 (Low) or 1 (High) µg/µL PB-pTetO-ZsG (tdTom -1.7±1.7; Low Ctrl 8.1±4.0; High Ctrl 140±20;) **E**,**F)** Scatter plots of ZsG intensity measured in all cells analysed from 0.1 (E, Low) or 1 (F, High) µg/µL PB-pTetO-ZsG electroporations. **G)** Lines of best fit for all conditions tested, based on scatter plots in E and F and for tdTom expressed alone. This demonstrates the robust induction and low leak of DOX inducible expression from high and low concentrations of electroporated PB-pTetO-ZsG/PB-pSyn-rtTA. (Line of best fit statistics presented as (slope, Spearman Correlation Coefficient): tdTom 0.0013, 0.45; Ctrl Low 0.0057, 0.30; Ctrl High 0.037, 0.52; DOX Low 0.20, 0.53; DOX High 1.11, 0.60; p<0.0001 for all Spearman Correlation tests). Scale 100 µm (B).

To account for varying efficiency of electroporation from animal to animal we normalized ZsG intensity to constitutively expressed tdTom and filtered out cells negative for ZsG but positive for tdTom (Figure S4, A-C, and see Methods). Quantification of ZsG:tdTom fluorescence ratios demonstrated that DOX induced highly significant expression of ZsG for both concentrations of PB-pTetO-ZsG delivered (Figure 2B-D). There was a significant statistical interaction between the effect of DOX and that of DNA dosage (2-way ANOVA, P<0.001), wherein normalized ZsG expression was significantly higher for 1µg/µL (High) PB-pTetO-ZsG than for 0.1µg/µL (Low) ZsG IUE (Figure 2C,D).

To assess possible leaky expression from the tetO, we first electroporated Cre under control of the tetO in confetti transgenic mice harbouring a Cre-dependent brainbow cassette. These animals yielded low-level recombination without DOX administration (Figure S5), demonstrating at least very low levels of leaky Cre expression. Low level leak of ZsG was also visible in subsets of cells in uninduced control animals electroporated with 1ug/uL PB-pTetO-ZsG (Figure 2B, white arrow heads). To quantify this, we compared animals electroporated with PB-ptTetO-ZsG that were left uninduced to animals electroporated with pCA-tdTom alone, to control for bleed-through from the red channel. Mean green levels in animals electroporated with low [PBpTetO-ZsG] (0.1µg/µL) IUEs, in the absence of DOX, did not differ significantly from those expressing tdTom alone (Figure 2E). High [PB-pTet-ZsG] (1µg/µL), on the other hand, showed faint leaky green fluorescence when compared to low [PB-pTetO-ZsG] (0.1 µg/µL) and tdTom alone (Figure 2E). However, that low level leak was only present in 25% of electroporated cells (Figure S4A,D). Taken together, these data demonstrate that incorporating the TetOn system into transposable elements circumvents silencing of the tetO during development in the IUE context, and that lowering the concentration of the inducible plasmid abolishes leaky expression in the absence of DOX.

### uTTOP Confers DOX-Mediated Gene Expression Across the Lifespan and in Other Cortical Areas

We next verified that uTTOP would overcome the silencing observed for episomal plasmids at P21. After inducing at P21, we observed robust induction of ZsG in layer 2/3 cortical neurons (Figure 3A). Given that we and others have observed episomal TetOn constructs to be active during development (i.e. P6) but silent in more mature neurons (Figure 1B), we wondered if using transposable constructs only delays silencing of the system. We therefore allowed animals to mature to an age of 20 months before induction. Although by this point the coelectroporated episomal tdTom expression conferred by pCA-tdTom had waned, we observed strong expression of ZsG with no apparent leak (Figure 3A).

**Figure 3.**
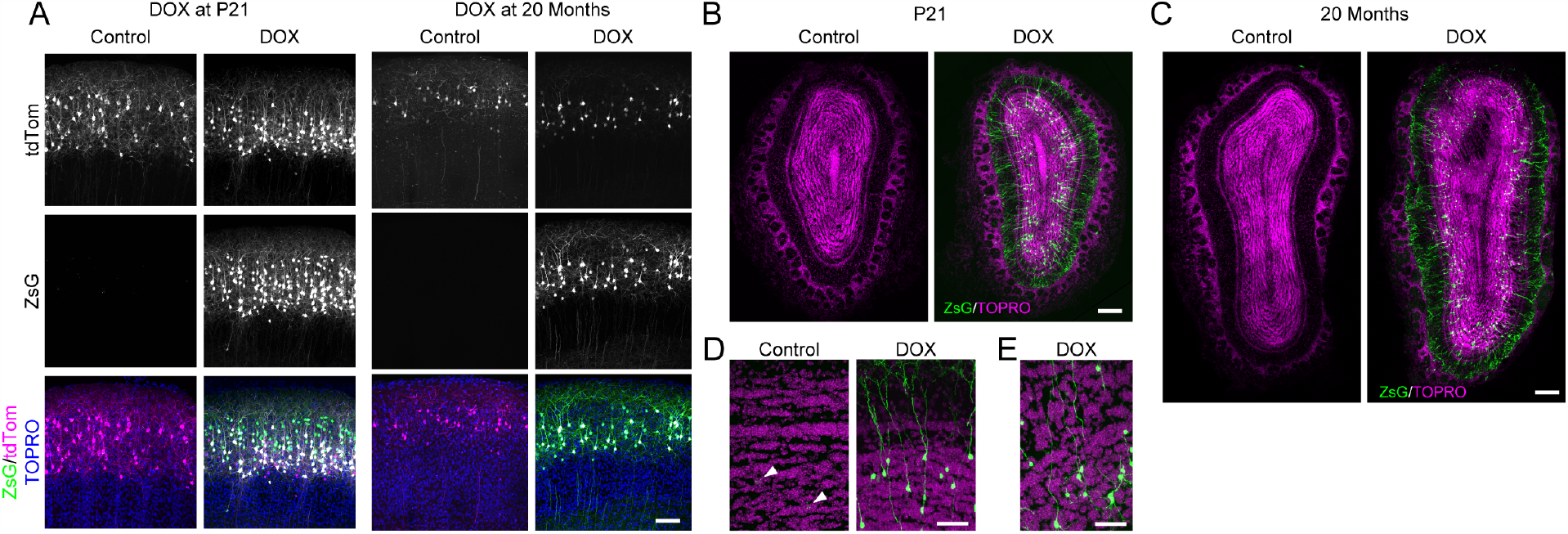
DOX-Induced ZsG Expresion by uTTOP at P21 and 20 Months in Neocortex and Olfactory Bulb. **A)** Expression of ZsG at P21 after 2 days of IP DOX administration (left), and at 20 months of age after 5 days of IP DOX administration (right). Weak tdTom expression was observed at 20 months compared to at P21 and P35. IUE of 1 µg/µL PB-pTetO-ZsG. Scale 100µm. **B**,**C)** DOX-induced expression of ZsG from transposable TetOn system in olfactory granule cells following induction at P21 (B) or 20 months (C) using the same induction protocols as mentioned in Figure 3. D,E) Higher magnification images showing ZsG in granule cells following induction at P21 (B) and 20 months (C). White arrow heads denote sporadic cells displaying low levels of leaky expression; no tdTom was observed and thus only the leak was able to indicate the presence of electroporated cells. At 20 months, no leak was visible and thus electroporated cells could not be identified. Scale 100µm (A) 200µm (B,C); 50µm (D,E)

We also wondered whether we could attain induction in the hippocampus and retrosplenial cortex following IUE of the uTTOP system to these areas with tripolar electrodes (dal Maschio et al., 2012; Szczurkowska et al., 2015). Clear DOX-mediated induction was observed at P35, but was only apparent in a subset of electroporated CA1 and subiculum neurons (Figure S6A). However, subsequent electroporations of PB-pTetO plasmids in other studies showed this is not a systematic issue (data not shown). In contrast, strong induction was observed in the retrosplenial cortex(Figure S6B).

### Olfactory Granule Cells Can be Targeted With uTTOP

Progenitors of the subventricular zone (SVZ) give rise to cortical neurons during development, as well as neural precursors that migrate through the rostral migratory stream to the olfactory bulbs throughout life (Takahashi et al., 2018). To investigate whether uTTOP plasmids stably integrate in the neurogenic niche of the SVZ that produces olfactory neurons, we examined the olfactory bulbs of animals electroporated with 1µg/µL PB-pTetO-ZsG, induced at P21 and 20 months. We found that at both time points ZsG expression was strongly induced (Figure 3B-E). Migrating precursors at the centre of the caudal extreme of the olfactory bulb also expressed ZsG in induced animals (Figure S7).

### Sonic Hedgehog Overexpression with uTTOP in Adult Cortical Neurons Upregulates Astrocytic Kir4.1

We previously showed that the morphogen, Sonic hedgehog (Shh), is expressed by mature neurons, and that the Shh signalling pathway is involved in regulating astrocytic molecular phenotypes (Farmer et al., 2016). However, it has yet to be shown whether Shh peptide itself is sufficient to change expression profiles of astrocytes **in vivo**. We therefore asked whether uTTOP could provide inducible expression with low enough leak to cause a Shh gain of function effect by promoting astrocytic expression of Kir4.1, a potassium channel whose expression is regulated by Shh signalling (Farmer et al., 2016) (Figure 2A). Shh undergoes posttranslational autocatalytic cleavage resulting in a non-signalling C-terminal fragment (Shh-C), and the fully mature N-terminal signalling peptide (Shh-Np, where “p” denotes “processed”). As native Shh is difficult to detect by immunofluorescence, an epitope-tagged recombinant N-terminal fragment (NShh-myc) is generally used in overexpression experiments. We first verified that the TetOn system was capable of producing secreted Shh signalling peptide following DOX induction (Figure 4B,C). Mature Shh-Np, which undergoes cholesterol modification as a result of its autocatalytic cleavage, retains more of its native signalling properties (Beug et al., 2011; Porter et al., 1996; Zeng et al., 2001). We therefore designed a construct (Shh-myc) coding for a full length Shh protein which, after undergoing auto-catalytic cleavage, leaves an epitope tagged autocatalytic domain as a reporter (Shh-C-myc), and a mature wild type signalling peptide (Shh-Np) (Figure 4D). We tested the ability of Shh-myc to drive Shh signalling in a luciferase reporter cell line (Gli-Luc C3H 10T1/2 cells) (Yam et al., 2009) and found that it performed as well as other constructs expressing wild type full length Shh and the N-terminal peptide alone (Figure 4F).

**Figure 4.**
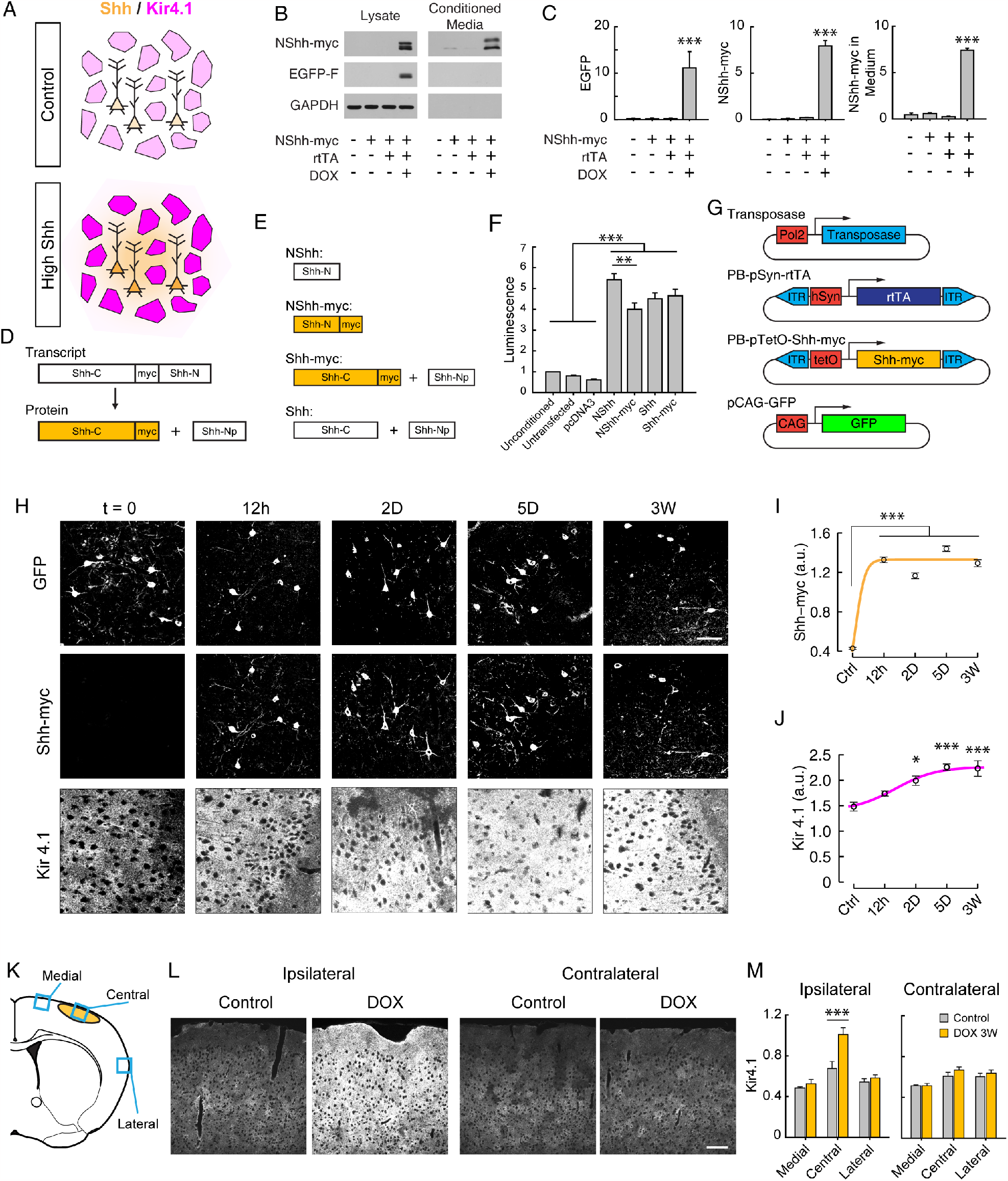
uTTOP-mediated expression of Shh in adult neurons drives astrocytic Kir4.1 expression. **A)** Schematic showing upregulation of astrocytic Kir4.1 by neuronal Shh. **B)** Western blots of DOX induced NShh-myc in HEK293T cells (left) and conditioned media removed from the same cell cultures (right). For following panels, GAPDH from cell lysates of the same wells that the medium was collected from was used as a loading control. Total protein from supernatant was used to adjust loading volumes. **C)** Summary plots corresponding to blots in A. (EGFP in a.u., untransfected 27.8±18.9; -rtTA,-DOX 72.4±40.8; -DOX 207.1±17.8; +DOX 7924.5±600.5, ANOVA p<0.001; NShh-myc in a.u., untransfected 159.5.8±66.9; -rtTA,-DOX 141.7±133.5; -DOX 159.4±105.7; +DOX 11093.3±3556.2, ANOVA p=0.002; NShh-myc in media in a.u., untransfected 444.7±199.6; -rtTA,-DOX 590.6±72.9; -DOX 236.9±59.0; +DOX 7405.4±220.7, ANOVA p=0.001). **D)** Schematic of transcript and autocleavage of the Shh-myc protein produced by PB-pTetO-Shh-myc. The mature signalling peptide is released, while the myc-tagged C-terminal autocatalytic domain is revealed by myc immunofluorescence. **E)** Schematic of protein products of panel of Shh expressing plasmids: pCAG-NShh (signalling peptide only) and pCAG-NShh-myc, pCAG-Shh-myc, pCAG-Shh (WT), respectively from top to bottom. Yellow fill indicates the peptides that can be easily detected with myc immunofluorescence. **F)** Summary plot of Shh signalling driven by expression of Shh peptides in (E) based on luminescence produced by C3H 10T1/2 cells stimulated with medium conditioned by HEK293T cells expressing indicated Shh peptides (Unconditioned 1; Untransfected 0.81±0.02; pcDNA3.1, 0.61±0.36; NShh 5.42±0.29; NShh-myc 4.00±0.31; Shh 4.51±0.28; ±18.9; Shh-myc 4.65±0.32; ANOVA p<0.001). G) Suite of plasmids, including transposable PB-pSyn-rtTA and PB-pTetO-Shh-myc, which were electroporated to test if Shh-myc overexpression in neurons can alter phenotypes of surrounding astrocytes. **H)** Exemplary images of Shh-myc induction, and corresponding Kir4.1 expression, after DOX administration timecourse. Images are shown in monochrome to clarify changes in fluorescence intensity. **I)** Quantification of Shh-myc expression timecourse (in a.u., Control 0.43±0.01, 12h 1.33±0.03, 2D 1.17±0.03, 5D 1.44±0.03, 3W 1.29±0.03, ANOVA p<2x10-16)J) Quantification of Kir4.1 expression in response to Shh-myc induction (in a.u., Control 1481.4±87.3, 12h 1739.3±51.6, 2D 1992.3±91.2,5D 2257.1±64.4, 3W 2231.3±152.1, ANOVA p=2.3x10-5).

**Figure 5.** **K)** Schematic of regions of coronal sections imaged to assess upregulation of Kir4.1 expression. Yellow denotes field of electroporated neurons. **L)** Comparison of Kir4.1 immunofluorescence in the centre of the electroporated fields (ipsilateral) and the same area in the contralateral hemisphere (contralateral). **M)** Quantification of Kir4.1 immunofluorescence in the regions shown in K in the ipsilateral (i.e. electroporated) hemisphere (Ipsilateral Central, in a.u., Ctrl 0.67±0.069, DOX 3W 1.01±.065; ANOVA p=0.005) and the same quantification performed in the contralateral (i.e. non-electroporated) hemisphere (left graph). Scale 50 µm (H), 100µm (L).

To test the ability of uTTOP to provide tight, conditional overexpression of Shh, cloned Shh-myc into a uTTOP plasmid (PB-pTetO-Shh-myc). We then coelectroporated this with PB-pSyn-rtTA and constitutively expressed EGFP in mouse somatosensory cortex (pCAG-GFP) (Fig. 4G). To assess induction kinetics of PB-pTetO-Shh-myc as well as the effect of Shh-myc induction, we performed a timecourse, administering DOX to P35 electroporated animals for 12 hours, 2 days, 5 days or 3 weeks (Figure 4H-J). Shh-myc expression peaked by 12 hours, indicating that uTTOP achieved rapid induction of gene expression (Figure 4I). Interestingly, Kir4.1 upregulation was slower, becoming signficant by 2D, and peaking at ∼5D days (Figure 4J). Kir4.1 expression levels can vary from brain region to brain region, and can also differ among neighboring astrocytes within a given brain region (Farmer and Murai, 2017; Tang et al., 2009). To take this into account in our analysis, we measured Kir4.1 immunofluorescence in the centre of the electroporated field in somatosensory cortex, medial to the electroporated field, and lateral to the electroporated field (Figure 4K), and found that Kir4.1 was significantly upregulated in the central field of induced animals, but not in uninduced animals (Figure 4L,M). Furthermore, Kir4.1 upregulation did not extend laterally or medially out of the electroporated field. The specificity of the effect was further confirmed by quantifying Kir4.1 expression in the contralateral hemisphere, which showed no change (Figure 4L,M). Taken together, these results show that neuronal Shh is sufficient modulate Kir4.1 expression in neighboring astrocytes. Thus, using uTTOP has allowed us untangle the spatio-temporal dynamics of neuron-astrocyte Shh signalling and to dissect the kinetics of non-cell autonomous changes to astrocyte properties over a period of several days.

## Discussion

Dissection of gene function, particularly during development and learning, requires fast and strong experimental induction of gene expression in the mouse brain. Here we examined the ability of a new method, uTTOP, to allow for fast DOX-mediated expression of a gene of interest. Although episomal TetOn plasmids provided adequate DOX-mediated induction in dissociated neurons and cortical neurons of young pups *in vivo*, these constructs were of no use in more mature animals where there was little to no induction. However, when the TetOn system was incorporated into transposable elements to create uTTOP, we observed robust DOX-mediated induction of gene expression across all time points examined and in multiple brain regions.

The method of choice for conditional gene expression is Cre-mediated recombination. Specific promoters driving Cre expression have been employed to control the timing of recombination (Lakso et al., 1992). For more precise temporal control, CreER and FLPeR approaches are often used (Luo et al., 2018; Matsuda and Cepko, 2007). However, CreER/FLPeR-based strategies can require long dosing periods with tamoxifen and provide variable degrees of recombination (Farmer et al., 2016; Slezak et al., 2007). Tamoxifen, an estrogen analog, is also known to alter dendritic spine numbers and morphology in both male and female rodents, introducing a potential confounding factor when using these systems in the brain (González-Burgos et al., 2012). Similar concerns apply to ligand-based split-Cre complementation, which requires rapamycin administration (Jullien et al., 2007). Photo-activatable recombinases have also been used for conditional expression in the brain, but require intrusive implantation of fibre optics (Kawano et al., 2016; Li et al., 2020; Meador et al., 2019). Destabilized-Cre stabilization has been demonstrated to allow for relatively fast expression in the brain, and has been used elegantly to dissect neuronal developmental questions (Pieraut et al., 2014), however, the lag to peak expression is 50-75h, leaving room for improvement (Sando et al., 2013). Here, we have demonstrated that uTTOP induces rapid, high levels of expression of a gene of interest in neurons in a variety of brain regions, with a lag time of 12 hours to peak expression. One concern when using binary conditional expression strategies that rely on TetOn/Off, as opposed to recombination techniques, is leaky expression from the tetO. TetOn systems have produced mixed results in terms of leaky expression in the absence of DOX. In early testing using transient transfection, rtTA was observed to drive low level but consistent basal expression from the tetO (Gossen and Bujard, 1992). However, stable cell lines had markedly lower leaky expression from the tetO (Gossen et al., 1995). Indeed, the few mouse lines that successfully implemented the TetOn system in the brain were not observed to have significant problems with leak (Baron and Bujard, 2000; Mayford et al., 1996; Yamamoto et al., 2003). Our results indicate that very low levels of leaky expression from the tetO does occur when using uTTOP with high concentrations of the inducible plasmid, but only in small subsets of electroporated neurons (Figure 2). Furthermore, by lowering the dose of PB-pTetO-ZsG delivered during IUE, leaky expression of ZsG was effectively abolished (Figure 2D). Our experiments with Shh (Figure 4) also demonstrate that the level of leak from uTTOP plasmids is negligible. Shh is an apt test case for assessing leakiness as it is well known to be a very potent morphogen, capable of functioning at very low concentrations (dynamic range >1nm) (Dessaud et al., 2010). This is particularly true when Shh is present in its native, processed form following autocatalytic cleavage, as is the case in our study (Beug et al., 2011; Zeng et al., 2001). Shh induction altered astrocytic Kir4.1 expression, while control tissue, as well as tissue from contralateral hemispheres of both control and Shh+ conditions, showed no change. Taken together, our work demonstrates that uTTOP provides tight conditional expression in neurons, with best-in-class induction kinetics.

It has been suggested that tTA-mediated transactivation is superior to rtTA-mediated transactivation due to its low levels of basal leak (Baron and Bujard, 2000). However, others highlight downsides of the tTA system – namely that tTA requires long-term, and in most cases in utero administration of DOX, which is also a broadspectrum antibiotic. Furthermore, even in cell culture, the TetOff system displays slow kinetics for optimal induction (up to 216h) (A-Mohammadi et al., 1997), and in mice, the build-up of the lipophilic DOX in bone and muscle results in a 15 to 20 day lag in induction once DOX administration is terminated (Furth et al., 1994; Kistner et al., 1996; Mansuy and Bujard, 2000; Mansuy et al., 1998; Mayford et al., 1996). Given these points and our desire for high temporal precision, we opted to risk the potential leakiness of rtTA in favor of its more optimal induction kinetics. (Indeed, herein lies a major benefit of harnessing IUE with uTTOP – riskier approaches that would otherwise be avoided when designing a transgenic animal can be employed more confidently when start-up only involves the minimal time and financial costs of basic cloning and as little as a single surgery, as opposed to creating a transgenic mouse model.) Nonetheless, we found that IUE of the TetOn system demonstrated very low levels of leaky expression, and that this can be effectively abolished by lowering the amount of inducible DNA electroporated. Varying the dose of electroporated rtTA may also help to address this. Finally, in cases where very low leak is still an issue, uTTOP can be combined with improved Tet operators (Loew et al., 2010) and post-transcriptional regulatory strategies (Zhou et al., 2020), both of which have been employed to reduce background expression of extremely toxic genes under TetOn control.

The use of piggyBAC transposase expression systems is also not without drawbacks. Notably, integrating transposable elements into the genome at TTAA sequences likely disrupts coding and non-coding sequences, and it should be noted the piggyBAC system displays a preference for targeting transcription units (Ding et al., 2005). While disruption of gene function by piggyBAC-mediated transposition cannot be neglected, the relatively random nature of the integration at the common TTAA nucleotide consensus sequence likely results in widely varying mutations in the population of electroporated cells. Thus, assessing phenotypes associated with the electroporated gene of interest across a large population of cells should control for deficits induced in individual cells, or small subsets of cells. Confounds associated with piggyBAC-associated mutagenesis can also be avoided with carefully designed negative controls. Furthermore, as opposed to other transposases, piggyBAC does not cause largescale genomic instability, and also displays a very low likelihood of leaving footprint mutations following excision due to persistent transposase activity (Yusa et al., 2011).

### Shh Overexpression Alters Astrocyte Molecular Phenotype

Previous work from our group and others has demonstrated that the Shh is expressed in neurons and that the Shh signalling pathway is involved in diversifying the molecular profile of neighboring astrocytes (Farmer et al., 2016). Driving this pathway via a constitutively active Shh receptor in astrocytes caused broad transcriptomic changes in cerebellar and cortical astrocytes. The current results provide the first direct evidence that the Shh peptide itself is capable of changing the expression profile of astrocytes in the adult mouse *in vivo*. Importantly, Shh binds its receptor, Patched, with high affinity (Stone et al., 1996), suggesting that appreciable leak from PB-pTetO-Shh would be detected. However, control animals did not show an elevation in Kir4.1 expression relative to adjacent or contralateral non-electroporated brain areas, suggesting that uTTOP does not display problematic levels of leaky expression in the absence of DOX.

### Future Directions

While we found that a dose of 50ug/g DOX provided good induction of gene expression (Figure 6F) it should be noted that this is a high systemic dosage of DOX. This likely leads to changes in the microbiome due to the antibiotic actions of DOX, and may also cause other adverse effects as DOX cytotoxicity has been reported in the context of Tet-regulated gene expression (Ermak et al., 2003). These issues may lead to altered behaviour, and indeed changes in the microbiome may alter brain function (Foster and Neufeld, 2013). It is therefore essential that rigorous controls are incorporated into the experimental design when applying uTTOP.

Potential off-target effects of DOX can also be mitigated by more advanced dosing strategies. Intravenous injections of DOX may allow for smaller and/or more concentrated doses. Furthermore, DOX has poor kinetics in crossing the blood brain barrier (Beard et al., 2006), with a cerebrospinal fluid availability only 15% of that in serum (Dotevall and Hagberg, 1989). It has been found that the tetracycline derivative, metacycline, is capable of driving expression of TetOn systems, and is less toxic at high doses (Krueger et al., 2004). Metacycline may therefore be useful in cases where greater and longer induction is necessary. Importantly, metacycline’s serum half-life is shorter than DOX, which may be either detrimental or beneficial if a faster run down of induction is desired. Alternatively, 9-tert-butyl-DOX (9TBDOX), a more lipid-soluble derivative of DOX, has been reported to have a 10-fold higher binding affinity for the wild type Tetracycline Repressor (Zhu et al., 2007). While these strategies will likely improve both adult inductions and inductions of pups through the mother’s milk, further improvements can also likely be achieved in pups via experimental design. DOX builds up in muscle and bone (Mansuy and Bujard, 2000). Taking this into account, foster mothers could be pre-loaded with DOX prior to birth of pups, and pups could be transferred to the foster mother at the time when induction is desired, ensuring a maximal dose of DOX in the milk as soon as fostering occurs. Finally, combinations of DOX delivery by injection, in the diet, and in the drinking water with appropriate supplements (Cawthorne et al., 2007), should be systematically tested to find the optimal dosing regimen for different induction needs.

Finally, proper titration of transactivator levels must also be considered. As can occur with Cre (Forni, 2006; Schmidt-Supprian and Rajewsky, 2007), excessive expression of rtTA can cause transcriptional squelching whereby exogenous transcription factors sequester endogenous transcription machinery to such an extent that cellular health is compromised (Baron and Bujard, 2000). IUE results in varying levels of expression throughout the field of electroporated neurons. While the variable efficiency of IUE on a cell-by-cell basis may be relied on to dilute out effects of squelching when considering a population of neurons, these effects can be controlled for by including a rtTA-lacking condition. However, it may be optimal to use rtTA-transgenic mice with moderate and consistent rtTA levels (e.g., Mansuy et al., 1998), electroporating only the inducible plasmid expressing the gene of interest. This approach also opens the possibility of using cell-type specific rtTA driver lines.

We have presented a novel combination of tools that allows for rapid-onset, robust expression of genes across the lifespan of the mouse in a variety of brain regions Taking the current results into consideration, along with possibilities for honing this approach detailed above, uTTOP represents a powerful new tool for conditional gene expression in neurons.

## Acknowledgements

The authors would like to acknowledge technical assistance from Dr Emma Jones, Michael Pratte, Nensi Alivodej, and Amy Zhou. We would like to thank Dr Min Fu and Shi-Bo Feng at the Molecular Imaging Platform of the Research Institute of the McGill University Health Centre for services provided, as well as Drs Andrew Greenhalgh, Edward Ruthazer and David Stellwagen for comments on experiments and the manuscript. The pCAG-Cre-myc vector was kindly provided by Dr Artur Kania, and in utero electroporation training was provided by Julie Cardin of the Kania Lab. The hSynapsin promoter was extracted from a pSKN-hSynapsin backbone kindly provided by Dr Ellis Cooper. The Gli-Luciferase reporter cell line was kindly provided by Dr Jorge Filmus by way of Dr Frédéric Charron. This work was supported by the Canadian Institutes of Health Research (MOP111152, MOP123390, and PJT156247) to K.K.M.

## Materials and Methods

### DNA Plasmids and Molecular Biology

pTetO-ZsGreen (pTetO-ZsG) was obtained from Clontech (pmRi-ZsGreen; Cat. No. 631121) and all other pTetO promoters are copies of the PTet-14 promoter (Urlinger et al., 2000), which matches that of pmRi-ZsGreen. The rtTA-expressing plasmids used in this study were constructed from the rtTA2^*S*^-M2 (Urlinger et al., 2000). All plasmids engineered for this study were verified by Sanger sequencing provided at the McGill and Genome Quebec Innovation Centre. Sequencing was performed with primers designed to attain full coverage on both strands of the coding regions of the genes of interest to ensure the absence of mutations. Novel plasmids engineered for this study are available on request.

### Animals and Animal Care

Experiments were approved by the Montreal General Hospital Facility Animal Care Committee and followed guidelines of the Canadian Council on Animal Care. The R26R Confetti Brainbow knockin mouse was obtained from The Jackson Laboratory (R26R Confetti, Stock Gt(ROSA)26Sortm1(CAG-Brainbow2.1)Cle/J, Stock No. 013731).

### *in utero* Electroporation

IUE was performed on or timed-pregnant ICR/CD1 mice ordered from Charles River Laboratories (Senneville, Québec). C57BL/6 mice (Figures S2,S5 only) were produced from an in-house colony. Mice were housed onsite for a minimum of 2 days before surgery. Mice were kept on a 12/12 light/dark cycle and allowed access to food and water ad libitum.

Pregnant mice were anaesthetized with 5% isoflurane and maintained with 2% isofluorane on a homeothermic warming pad for the duration of the IUE surgery. Opthalamic ointment was used to prevent drying of the eyes. Breathing, temperature and color of mucosal membranes were monitored throughout the surgery. If any aberrations were observed, isoflurane flow was temporarily decreased until symptoms were resolved. The abdomen was shaved and cleaned with providone and isopropanol wipes. A 1.5cm vertical incision was then made at the midline of the abdomen with scissors. Sterile gauze (soaked with sterile, 39°C phosphate buffered saline (PBS)) was positioned around the incision to provide a clean, well hydrated surface on which to place the uterine horns. The uterine horns were extracted from the peritoneal cavity by massaging the abdomen with PBS-wetted gloves, and laid on the moist gauze. Uterine horns were kept moist and warm by frequent application of warm PBS throughout the surgery. A pulled glass micropipette attached to a foot pedal-controlled microinjector (Harvard Apparatus, MA) was used to deliver 2µL (E13-E14) or 3µL (E15E16) of 2-3.5µg/µL solution of DNA to the lateral ventricle of the embryo’s brain. Micropipettes were bevelled at a 35-45°angle using diamond lapping film attached to a repurposed computer hard drive. The bevelled, sharpened pipet tip helped to minimize damage to the uterine lining and the CNS. DNA was dissolved in endotoxin free water or TE buffer and brought to the final concentration in PBS and 0.03% Fast Green dye. Five to seven 36-42V pulses of approximately 50ms were applied horizontally across embryo’s cranium using an BTX ECM 830 square pulse generator and 3mm platinum-coated tweezer electrodes (BTX, MA). The embryos on either side of the uterine corpus were left untouched, otherwise all embryos were electroporated unless excessive manipulation was required to expose the top of the cranium. The incision was then flushed with warm PBS, the abdomen was closed with 6-0 nylon running sutures, and the skin with 5-0 nylon running sutures. The mother was then returned to its home cage and allowed to recover until mobile and grooming resumed in a 28°C recovery chamber. Wet food and carprofen analgesia were provided following the surgery.

All IUEs were performed at E15-16, except for hippocampal IUEs which were performed at E13-14.

### Perfusion and Tissue Preparation for Light Microscopy

Electroporated mice (6 months old; male and female; CD1) were anesthetized with isoflurane and tissue fixed by transcardial perfusion first with ice cold DPBS followed by 4% paraformaldehyde in 0.1 M phosphate buffer (pH 7.4) at 5 mL/min for 5 minutes. Brains were removed and post-fixed in the same solution overnight at 4°C. Following equilibration in 30% sucrose in DPBS, the tissue was embedded in OCT (Somagen, Edmonton, AB) and snap frozen. Brains were cryosectioned coronally at 30 µm thickness and stored in DPBS at 4°C until use. Sections were mounted onto charged Superfrost Plus glass slides and preserved with SlowFade Gold Antifade Mountant (Thermo Fisher, Waltham, MA). Confocal microscopy was performed using an Olympus FV1000 laser scanning confocal microscope with a 60x oil-immersion objective (N.A. 1.4) at a resolution of 0.132 µm^2^/pixel. Stuctured Illumination Microscopy (SIM) image stacks were acquired using a Zeiss LSM880 ElyraPS1 microscope with a 63x oil-immersion objective (Plan-Apochromat 63x/1.40 Oil DIC). All images were obtained using 5 rotations and processed for SIM using the Zeiss Zen software.

### Doxycycline Administration

100µg/g DOX per day was administered in PBS solution via IP injection by two doses of 50µg/g (5 mg/mL). Induction of hippocampal expression was performed with one 50µg/g dose per day. For 20 month-old mice, DOX was administered in a malodextrine based diet containing 3mg/g DOX (Custom Diet #TD.170534, Envigo) and once daily IP injections (50µg/µL) for 5 days. For 3-week induction of Shh-myc expression from PB-pTetO-Shh-myc, mice were administered DOX diet ad libitum. Fresh DOX diet was provided every 3 days.

### Tissue and Cell Culture

Primary hippocampal neuronal cultures were prepared as described previously in detail (Jones et al., 2012). Briefly, hippocampi were extracted from postnatal day 0 C57BL/6 mice, incubated in 0.1% papain, 0.02% BSA in Neurobasal-A medium (NBA) (Invitrogen) at 37°C for 15 minutes with periodic agitation. Hippocampal tissue was then transferred to warm NBA containing 1% egg white trypsin inhibitor and 1% BSA and triturated 8 to 10 times with fire polished Pasteur pipettes of decreasing pore diameter, discarding undissociated debris after each trituration. Neurons were then plated on coverslips coated with poly-L-lysine (0.1mg/mL) at 40,000 cells/well in 500µL of neuronal culture medium in 24 well plates. Neuronal culture medium contained NBA supplemented with 2% B27 (Invitrogen), 1mM GlutaMax (Invitrogen) and 1% penicillin/streptomycin, and one third of medium was changed every 2 to 3 days by removing 200µL and adding 300µL. Cytarabine (ara-C) was added to a final concentration of 3µM after 3 days *in vitro* to prevent glial overgrowth. Neuronal transfections were performed with lipofectaine at 6 to 8 days *in vitro*. For each well, 75µL of unsupplemented NBA containing 1 to 2µg DNA was combined with 75µL unsupplemented NBA containing 3µL of lipofectamine. While this mixture was allowed to stand for 20 to 40 minutes, neurons were transferred into 400µL of equilibrated unsupplemented NBA in 24 well plates. Transfection mixtures were then added to neurons dropwise and incubated for 2 to 3 hours before being transferred back to their conditioned neuronal culture medium. Expression of transfected plasmids proceeded overnight, followed by DOX-induction for 24 to 48h.

HEK293T and 8XGli-Luciferase C3H 10T1/2 (Gli-Luc reporter) cells were cultured in medium containing 10% FBS/ 1% Penicillin-Streptomycin/DMEM and passaged at a ratio of 1:10 when 70-90% confluent.

### Immunofluorescence

Mice were transcardially perfused with PBS wash followed by 4% paraformaldehyde/0.1M phosphate buffer (PB), pH 7.4 (P8: 2mL wash, 15mL paraformaldehyde at a rate of 2mL per minute; P23: 3mL wash, 20mL paraformaldehyde, 3 mL/min; P35: 5mL wash, 35-40mL paraformaldehyde, 5mL/min; 20months; 5 mL wash, 50 mL paraformaldehyde, 5mL/min). Brains were post-fixed in 4% paraformaldehyde/0.1M PB overnight, cryoprotected in 30% sucrose/PBS, and embedded in OCT embedding medium. Free-floating coronal sections (40µm) of the entire cerebrum were cut, collected in PBS, and screened to locate the electroporation epicentre. Sections were permeabilized for 15 min in 1%Triton X-100 (TX-100)/PBS, blocked in 10% normal donkey serum (NDS) (Jackson Laboratories)/0.2% TrX100/PBS for 1.5-2h and stained with primary antibodies in 1% NDS/ 0.2% TX-100/PBS for 16 to 72 hours. Sections were then washed 3 times in 1%NDS/ 0.2% TX-100/PBS, incubated with fluorescent secondary antibodies (Alexa Fluors, Invitrogen) for 2 hours, washed 3 times in PBS and mounted in Slowfade Gold mounting medium (Invitrogen). Primary antibodies: mouse anti-mCherry (Clontech, Cat. No. 632543), chicken antiGFP (Abcam, ab13970), and mouse anti-myc (Santa Cruz, 9E10). TO-PRO-3 Iodide (TOPRO) (ThermoFisher) was used as a nuclear counter-stain.

Primary neuronal cultures were washed briefly with 4°C PBS and then fixed with a solution of 4% formaldehyde/4% sucrose/0.1M PB cooled to 4°C, left for 15 min at room temperature, washed three times with PBS, permeabilized in 0.2% TX-100/PBS for 15 minutes, and blocked in 10% NDS/PBS for 1.5 to 2h. Coverslips were incubated in primary antibodies with 1% NDS/PBS overnight at 4°C, washed 3 times in 1%NDS/PBS, incubated with secondary antibodies (Alexa Fluors) for 2 hours in 1%NDS/PBS, washed 3 times in PBS and mounted in Slowfade Gold. Antibodies: rabbit anti-γ2 (Alomone Labs, AGA-005) and chicken anti-GFP (Abcam).

### Microscopy and Image Analysis

All single frame example images presented in figures were acquired on an FV-1000 laser scanning confocal microscope (Olympus). Images of primary neuronal cultures were acquired using a custom Ultraview spinning disc confocal system (Perkin Elmer) attached to a Nikon TE-2000 microscope. Mosaics presented in Figures 3, S2, S3, S6 and S7 were acquired on an LSM780-NLO laser scanning confocal (Zeiss) and an Axio Observer Z1 spinning disk microscope (Zeiss). For analysis of DOX-mediated ZsGreen (ZsG) induction, we used an XLUMPlanFL N, 1.00 NA 20X objective mounted on an FV1200MPE laser scanning system equipped with a variable bandpass filter (Olympus). To optimally separate tdTomato (tdTom) and ZsG signals, we acquired fluorescent bands of 500-540 nm (ZsG) and 565-665 nm (tdTom). Fluorescence intensity in all cases was measured with corrected total cell fluorescence (CTCF = Sum of pixel values in ROI – [mean background pixel value * area of ROI]).

Both 0.1 and 1ug/uL PB-pTetO-ZsG produced a small population of ZsG-/tdTom+ cells. To better assess the magnitude of induction of the uTTOP system, we focussed on the ZsG+ cells, filtering out non-expressing cells using bleed-through of tdTom into the green channel to set a threshold of inclusion equal to the mean plus three times the standard deviation (12.24 a.u.) of green fluorescence from tdTom alone. This resulted in the exclusion of only 0.7 ± .04% and 0.1 ± 0.02% of the cells from the 1 and 0.1 µg/uL conditions, respectively (see Figure S4 A,B,D).

### Western Blot Analysis

Following 24h of induction with 1µM DOX, HEK293T cells were briefly washed with ice cold PBS and then lifted from wells with cell scraper in Triton Lysis Buffer (TLB) containing 20mM Tris ph7.4, 137mM NaCl, 2mM EDTA, 1% TX-100, 0.1% SDS, 10% glycerol, and supplemented with protease inhibitors and sodium orthovanadate. Cells were lysed on ice for 10 minutes with periodic agitation, and lysates were centrifuged at 16,000 rpm for 10 min. Supernatants were stored at -80°C in sample buffer containing 6% SDS and 16% β-mercaptoethanol. To assess NShh-myc expression in cell culture medium, medium was collected, supplemented with protease inhibitors, spun down at 16,000 rpm to remove any cells in suspension, and stored at -80°C in sample buffer. Standard SDS-PAGE Western blotting procedures were used. Immunoblotting was performed with anti-myc (Cell Signalling, 9B11, 1:10,000 in 5% non-fat milk), anti-GFP (Clontech, JL-8, 1:10,000 in 5% non-fat milk) and anti-GAPDH (Millipore, MAB374, 1:300,000 in 3% BSA).

### Luciferase Assay

C3H10T1/2 cells stably transfected with a construct containing 8 repeats of a minimal Gli promoter driving expression of firefly luciferase (8XGli-luc reporter) were plated at a density of 75,000 cells per well in 24 well plates at time (t) = 0 hours. HEK293T cells were plated at the same time at 100,000 cells per well (t = 0h). Cells were allowed to grow overnight. HEK293T cells were transfected with molar equivalents (1.5x1026 mol/well) of Shh constructs for 4 hours (start at t=16h) with Lipofectamine2000 (Invitrogen) as per manufacturer instructions, and expression was allowed to proceed overnight in 0.5mL of medium to concentrate secreted Shh peptide. Gli-luc C3H 10T1/2 cells were starved overnight in 0.5% FBS/DMEM (starting at t = 20h). At t = 36h, conditioned medium form HEK293T cells was centrifuged at 5000 rmp to remove any Shh plasmid-expressing cells from suspension, and conditioned supernatant was added to starved Gli-luc C3H 10T1/2 cells. After 24 hours (t = 60h), Gli-luc C3H 10T1/2 reporter cells were rinsed with cold PBS, lysed on ice in Luciferase Cell Culture Lysis Reagent (Promega) by scraping with the back of 200µL pipette tips (using a different tip for each well). Luciferase expression was assessed using a beetle luciferin-based assay system (Promega, Cat. No. PR-E1500), and luciferase activity was quantified with an LMax Luminometer (Molecular Devices). Briefly, 100µL of Luciferase Assay Substrate solution was added to 20µL of lysates, one well at a time, in a 96 well plate, followed by a 3s pause and a 10s read.

### Statistical Analysis

Data are presented as mean ± SEM. Statistical analysis was performed with SigmaPlot software. For mean comparisons, *p<0.05, **p<0.01, ***p<0.001. Student’s t-test and One and Two-way ANOVAs with Tukey HSD pariwise comparisons, preceded by a Shapiro-Wilks normality test, were used unless otherwise noted. For IUE, statistics were performed on datasets comprising cells analyzed from 3 to 6 animals, 2-3 coronal sections per animal, with the exception of Figure 6 F and G where the Ctrl and DOX 2D conditions comprised 2 animals each.

## Supplementary Information

**Figure S1.**
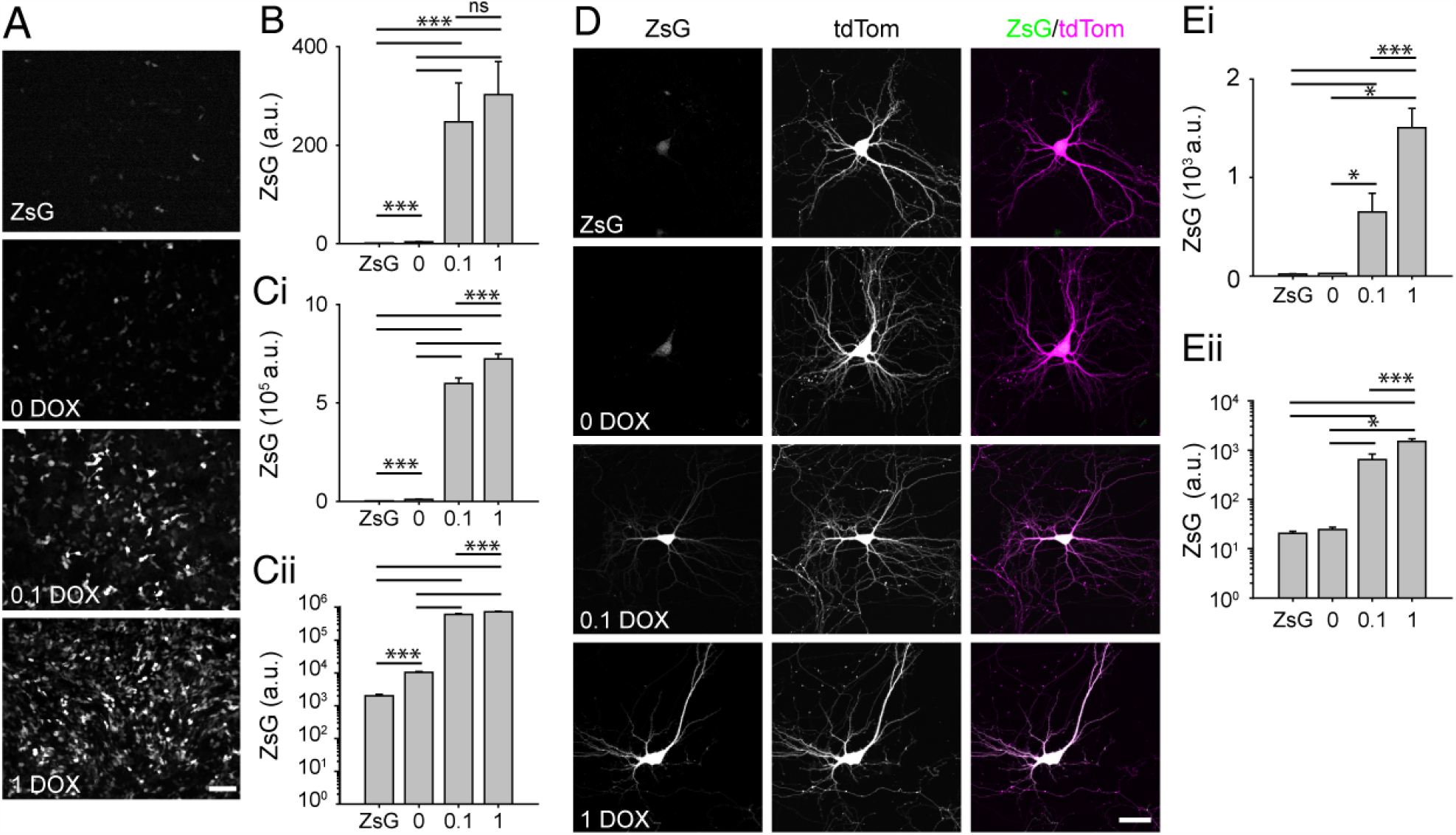
Episomal TetOn allows for conditional ZsGreen expression in HEK293T cells and cultured hippocampal neurons. A)DOX-induced ZsG in HEK293T cells, showing expression of ZsG when pTetO-ZsG was transfected alone (ZsG) or with pCMV-rtTA (with 0, 0.1 or 1 µg/µL DOX). B) Summary data of full frame ZsG fluorescence of near confluent cells (mean pixel value in a.u., ZsG Alone 1.36±0.075;0µg/µL DOX 3.46±0.87; 0.1µg/µL DOX 247.38±79.06; 1µg/µL DOX 302.75±67.16). C) Summary data of ZsG fluorescence in individual cells on a linear (Ci) and logarithmic scale (Cii). (CTCF in 103a.u., ZsG Alone 2.00±0.19 ;0µg/µL DOX 10.19±0.78; 0.1µg/µL DOX 598.33±28.73; 1µg/µL DOX 722.81±26.12; ANOVA p=0.002.) D) Dissociated hippocampal neurons cotransfected with pCA-tdTom and pTetO-ZsG alone (ZsG) or ZsG plus pSyn-rtTA (with 0, 0.1 or 1 µg/µL DOX). E) Summary data of ZsG fluorescence in individual neurons on a linear (Ei) and logarithmic scale (Eii). (In a.u., ZsG Alone 0.02±0.002 ;0µg/µL DOX .025±0.003; 0.1µg/µL DOX 0.65±0.19; 1µg/µL DOX 1.50±0.20; ANOVA p<0.001)

**Figure S2.**
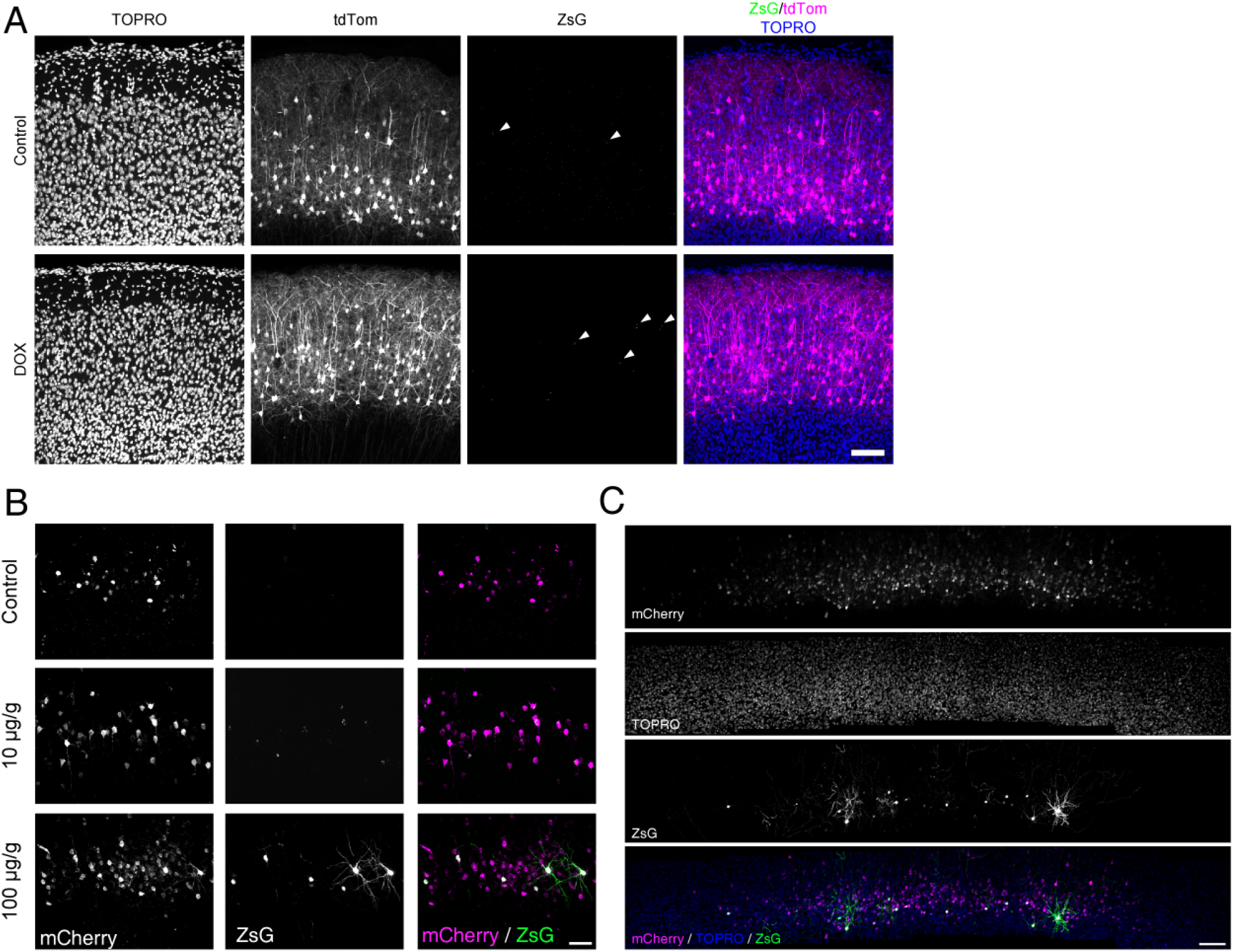
Absent or nearly absent DOX induction at P21 in CD1 and C57BL/6 animals, respectively. Example images demonstrating an almost complete lack of induction of episomal 0.1 µg/µL pmRi-ZsG at P21 after 100µg/g DOX per day for 3 days IP. White arrow heads denote cells displaying very low levels (a few weak puncta per cell) of ZsG expression in CD1 mice. B) Example images demonstrating sparse expression of episomal pmRi-ZsG (1 µg/µL) in C57BL/6 mice (10 µg/g DOX middle row, 100 µg/g DOX bottom row) at P21. C) Mosaic of a coronal section showing the most dense induction of ZsG at the rostrocaudal epicentre of the electroporated field in a C57BL/6 mouse (1ug/mL pmRi-ZsG, 100ug/g DOX for 3 days IP). Scale 100µm (A,C); 50µm (B).

**Figure S3.**
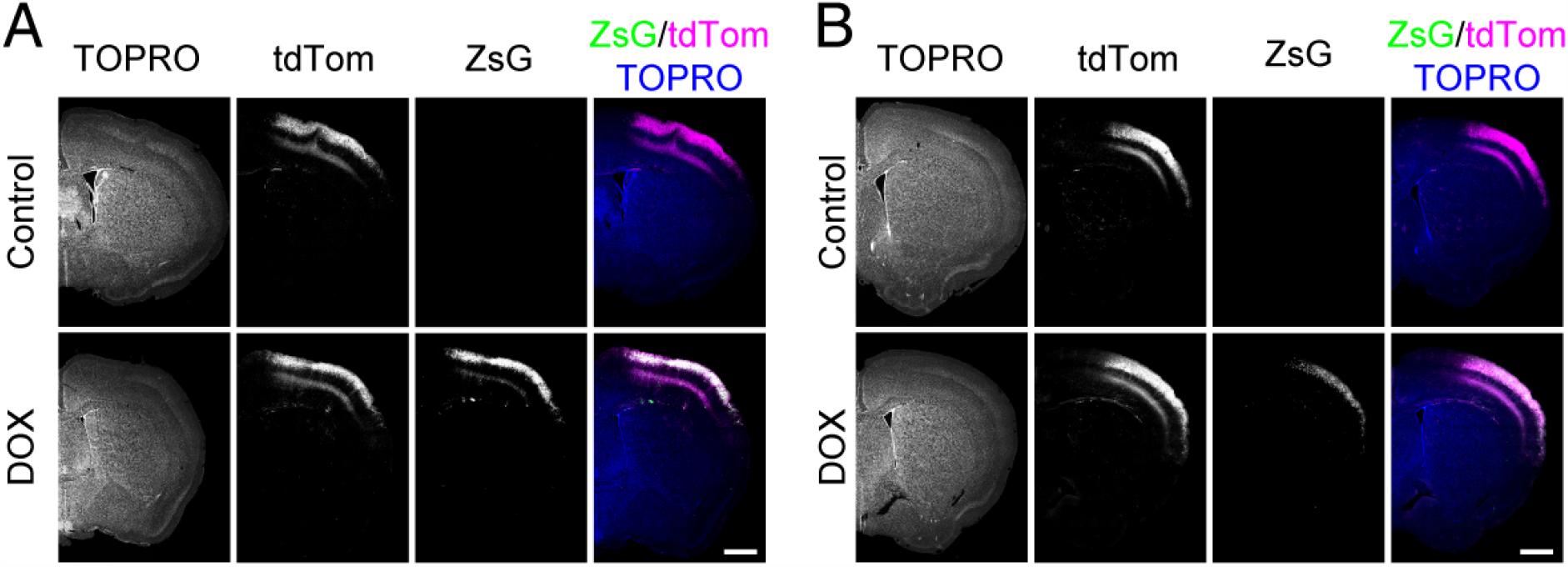
PB-TetO-ZsG induction across entire electroporated field. A,B) Images of whole hemispheres of mouse coronal sections showing induction of high levels of ZsG with 1µg/µL electroporation of PB-pTetO-ZsG (A), and lower levels of ZsG with 0.1µg/µL electroporation of PB-pTetO-ZsG (B). pCA-tdTom was used as a positive electroporation control, and TOPRO as a nuclear counter strain. Induction was 100ug/uL DOX for 3 days in both cases. Scale 500µm (A,B).

**Figure S4.**
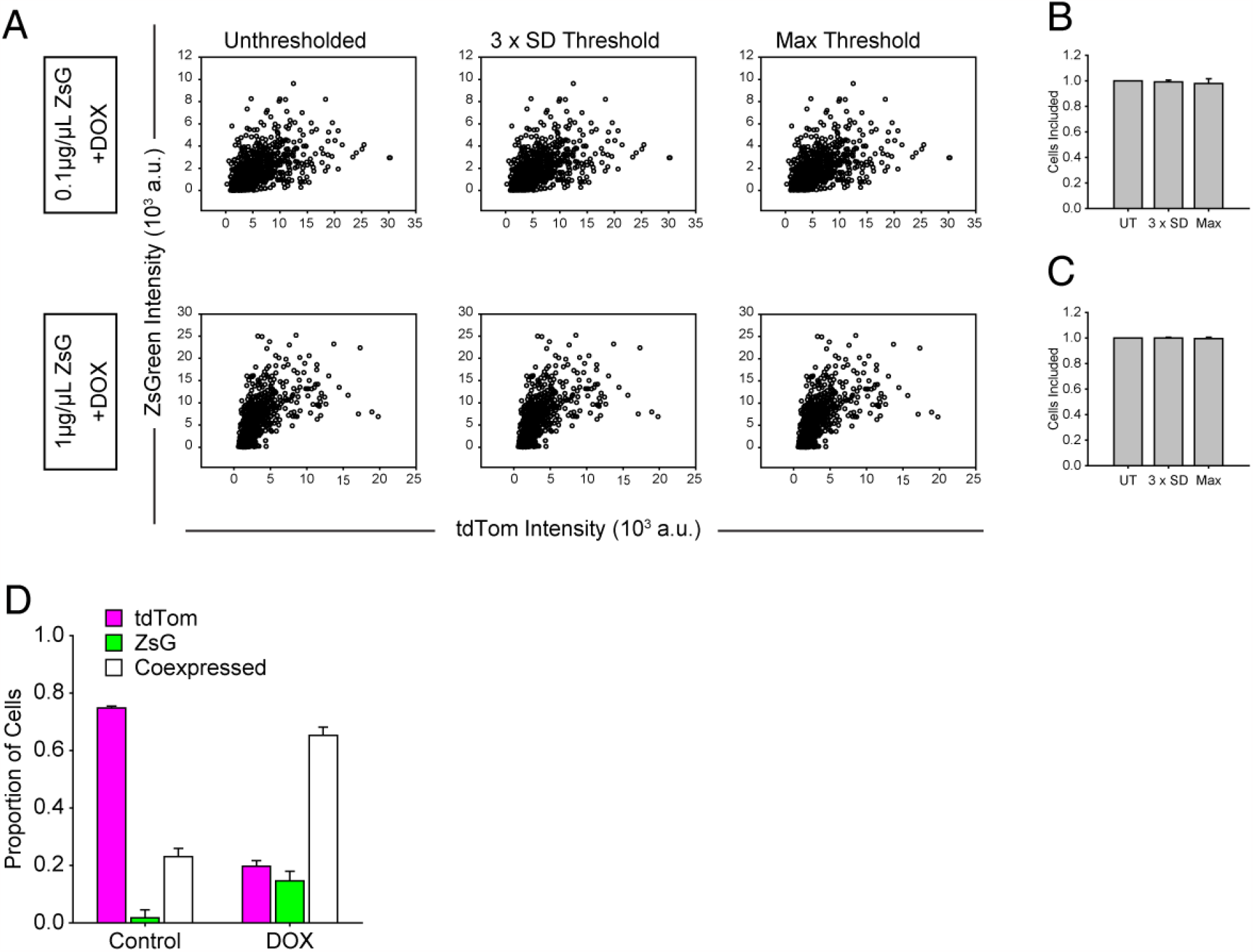
Proportions of thresholded neurons and neurons coexpressing tdTom and ZsG. A) Scatter plots of unthresholded data and with the two thresholds applied. B,C) Summary plots of the proportions of cells kept after thresholds were applied averaged across sections. UT=unthresholed, 3xSD = 3 x standard deviation, max = 29.9 a.u. (B: DOX Low, UT 1, 3xSD 0.99±0.004, Max 0.98±0.01; C: DOX High UT 1, 3xSD 0.998±0.002, Max 0.995±0.003) D) Proportions of cells expressing tdTom, ZsG or both in induced and uninduced animals that had been electroporated with 1 µg/g PB-pTetO-ZsG. (D, Ctrl 0.75±0.006 tdTom+, 0.019±0.03 ZsG+, 0.23±0.02 Coexpressed; G, DOX 0.20±0.02 tdTom+, 0.14±0.03 ZsG+, 0.65±0.03 Coexpressed.)

**Figure S5.**
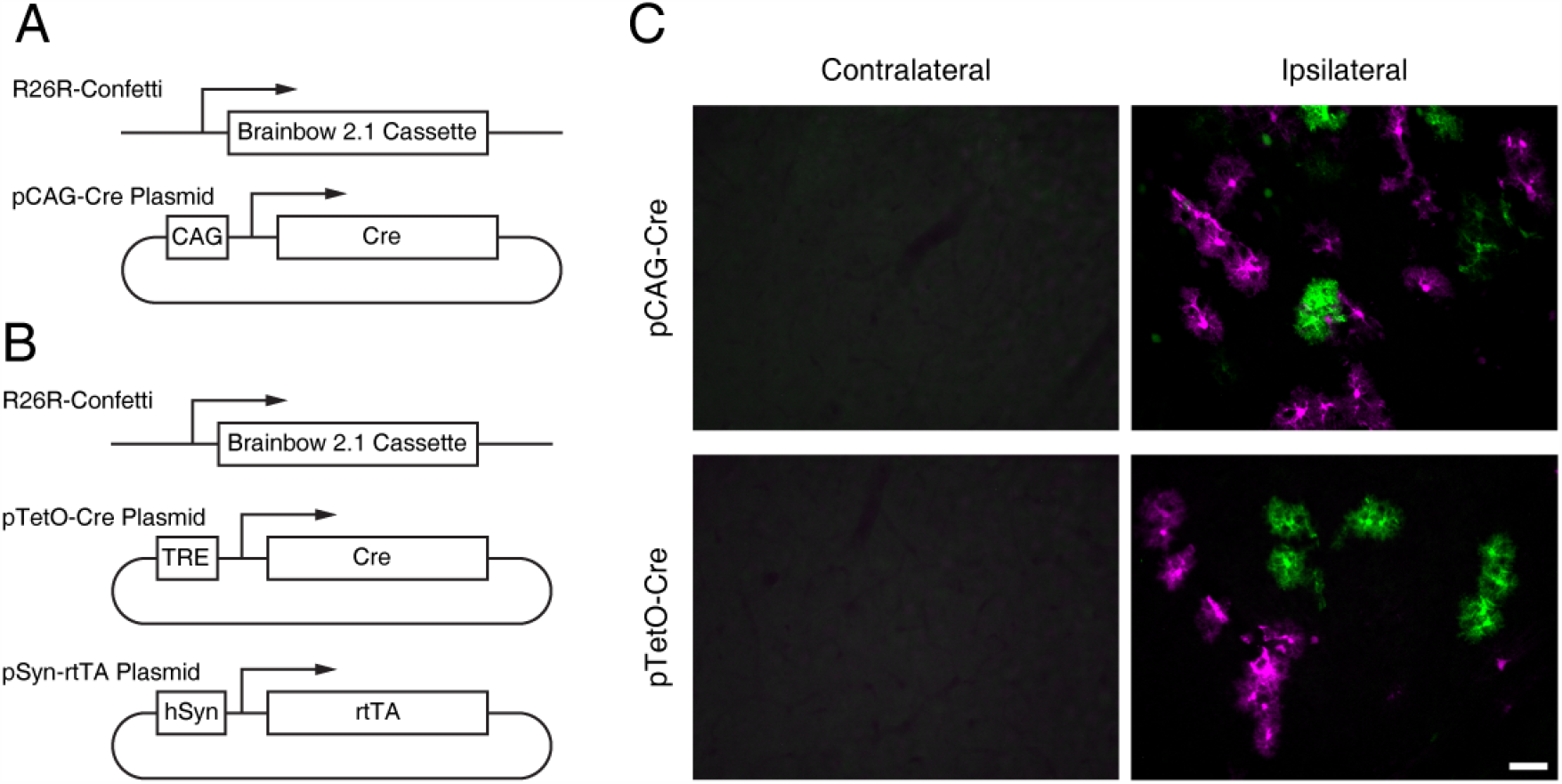
Leaky expression of Cre from episomal pTetO-Cre. Schematic of genomic Brainbow allele and electroporated plasmids for constitutively expressed Cre (A) and DOX-inducible Cre (B). C) Example image demonstrating that both sets of electrorporated plasmids expressed sufficient levels of Cre to cause recombination. Note that this mouse line is known to have higher expression in astrocytes and astrocytic labelling is thus much brighter than that of neurons. Scale 50µm.

**Figure S6.**
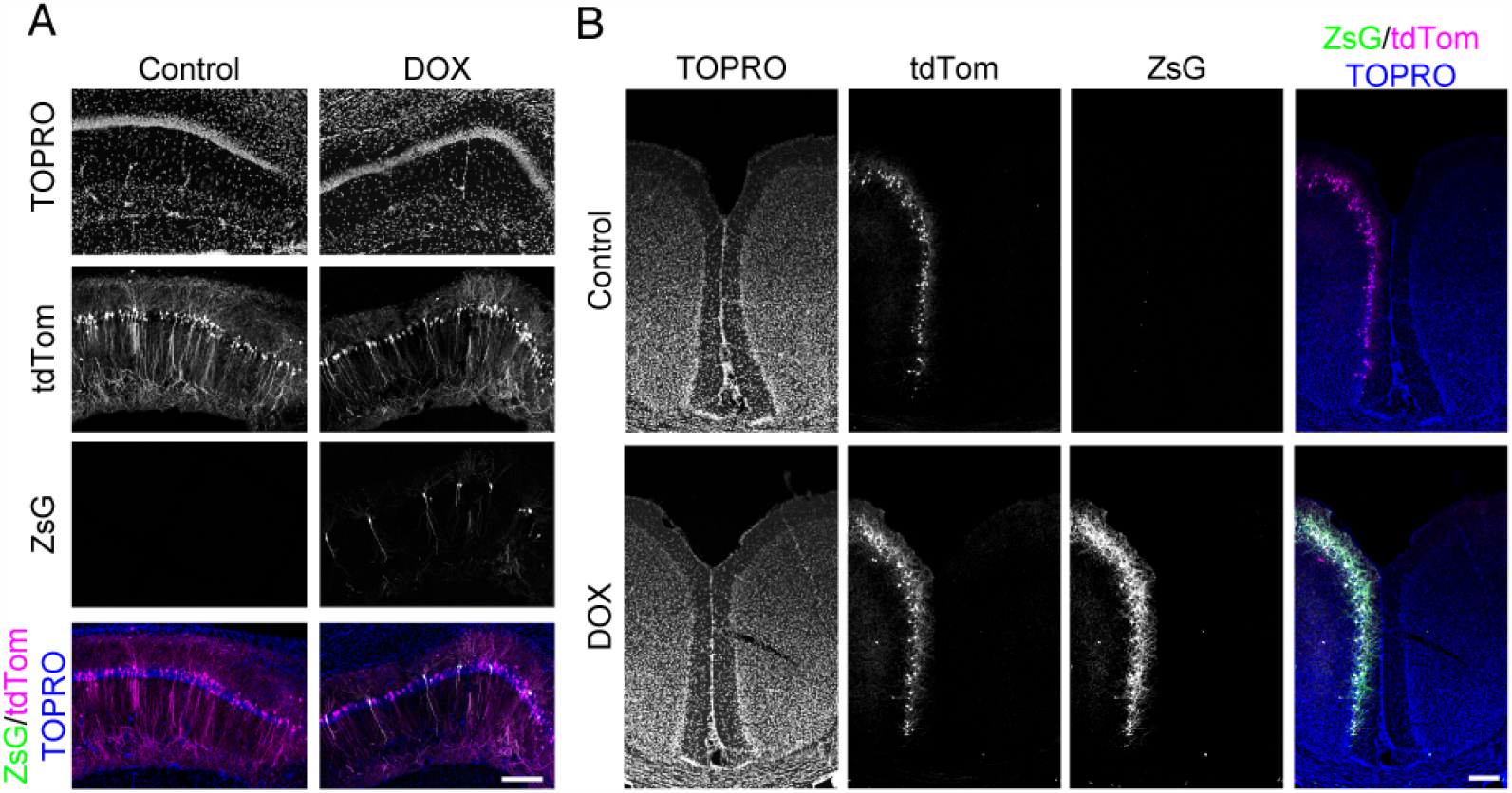
DOX induced expression in adult hippocampus and retrosplenial cortex. A) DOX induced ZsG expression in CA1 pyramidal cells of the hippocampus. B) DOX induced ZsG expression in retrosplenial cortex. Electroporation of 1 µg/µL PB-pTetO-ZsG, 100µg/µL DOX for 3 days at P35 in both A and B. pCA-tdTom was used as a positive electroporation control, and TOPRO as a nuclear counterstain. Scale 200µm (A), 100µm (B).

**Figure S7.**
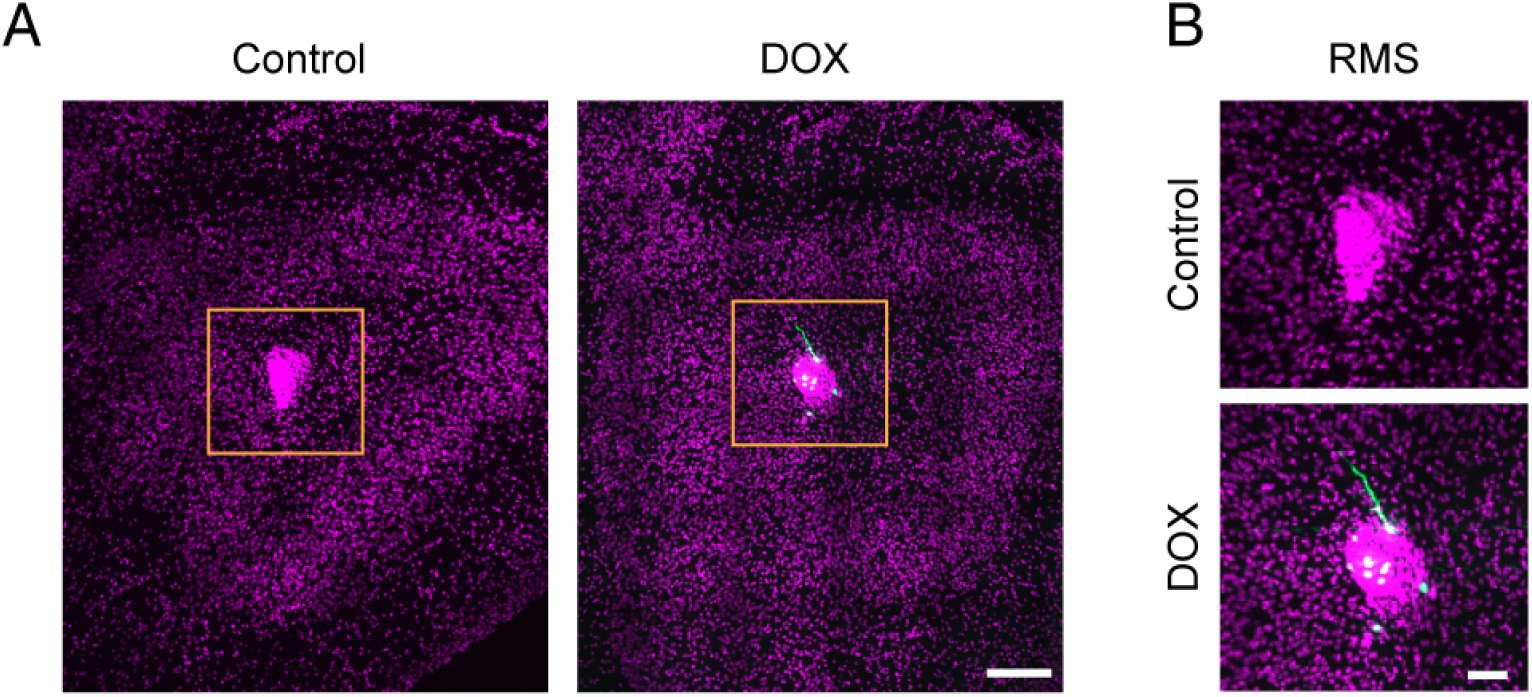
Induction of ZsG at P35 at caudal pole of olfactory bulb. A) Images of the caudal poles of the olfactory bulbs showing DOX-mediated ZsG expression in migrating precursors. B) Zooms of boxed regions in A. Scale 150µm (A); 50µm (B)

